# Activity of DNA polymerase κ across the genome in human fibroblasts

**DOI:** 10.1101/2024.02.22.581598

**Authors:** Mariela C. Torres, Dongxiao Sun, Thomas E. Spratt

## Abstract

DNA polymerase κ (Pol κ) is a specialized polymerase that has multiple cellular roles such as translesion DNA synthesis, replicating repetitive sequences, and nucleotide excision repair. We have developed a method for capturing DNA synthesized by Pol κ utilizing a Pol κ - specific substrate, *N*^2^-(4-ethynylbenzyl)-2′-deoxyguanosine (EBndG). After shearing of the DNA into 200-500bp lengths, the EBndG-containing DNA was covalently bound to biotin using the Cu(I)- catalyzed alkyne–azide cycloaddition reaction, and isolated with streptavidin beads. Isolated DNA was then ligated to adaptors, followed by PCR amplification and next-generation sequencing (NGS) to generate genome-wide repair maps. We have termed this method polymerase κ sequencing (polK-seq). Here we present the human genome maps for pol κ activity in an undamaged cell line. We found that pol κ activity was enhanced in euchromatin regions, the promoter of genes, and in DNA that is replicated early in S-phase.

## Introduction

DNA polymerase κ (pol κ) is one of the sixteen DNA-dependent DNA polymerases expressed in humans (1). Pol κ is typically described as a translesion DNA synthesis polymerase because of its exceptional ability to bypass bulky *N*^2^-alkyl-dG adducts (2, 3). But pol κ has been linked with nucleotide excision repair (4), cross-link repair (5), the ATR activation pathway (6), and replication of repetitive DNA sequences (7). To examine the activity of pol κ in the genome in an unbiased manner, we utilized the *N*^2^-4-ethynylbenzyl-2’-deoxyguanosine (EBndG, Figure 1a), a nucleoside that has high selectivity with pol κ. Thus, DNA that had been synthesized with pol κ was captured using the Click reaction and the capture DNA mapped to the human genome. We found that pol κ is active on euchromatin and during early S-phase.

**Figure 1.**
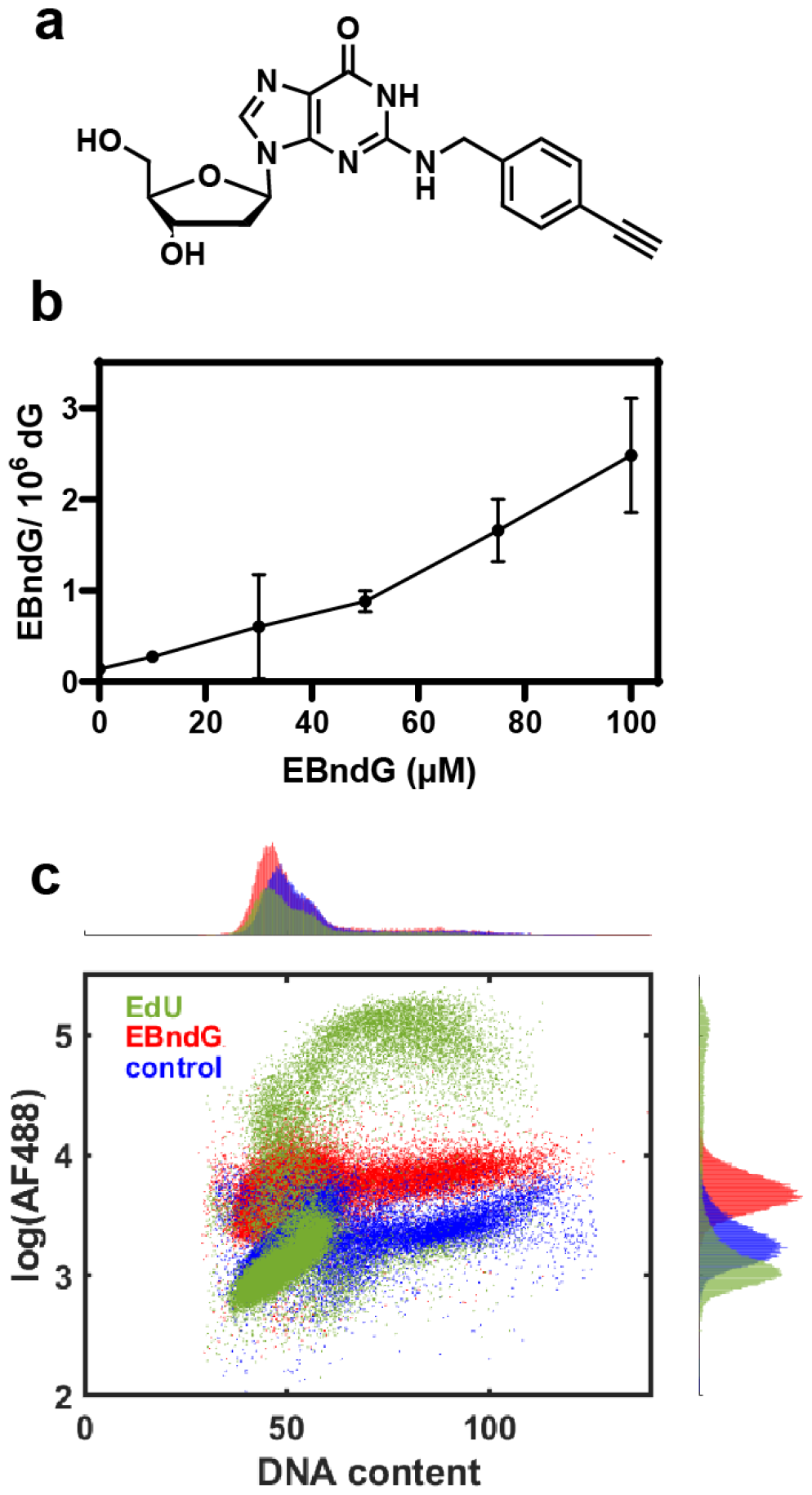
Incorporation of EBndG into the genome. (a) EBndG structure. (b) Incorporation of EBndG into the DNA of GM12878 cells at different concentrations of EBndG. Data are the mean of two experiments (± SD). (c). Flow cytometric analysis of the incorporation of EdU (50 μM) and EBndG (100 μM) into GM12878 cells over 4 h.

## Results

### EBndG incorporation

DNA, isolated from GM12878 cells treated with EBndG for 4 h, was enzymatically hydrolyzed to nucleosides. The level of EBndG was analyzed by monitoring the neutral loss of deoxyribose by HPLC-MS/MS. The level of EBndG incorporated was proportional to the concentration and is shown in Figure 1b. The incorporation was also investigated using flow cytometry. The results in Figure 1B, show the differences in incorporation between EdU and EBndG. As expected, EdU was extensively incorporated during S-phase. In contrast, EBndG was incorporated into the DNA at a much lower level, but the incorporation appears to be during all phases of the cell cycle.

### PolK-seq

GM12878 cells were treated with 0, 20 or 100 μM EBndG. After 4 h, genomic DNA was isolated, fragmented and DNA strands containing EBndG were covalently bound to Dde biotin picolyl azide with the Cu(I)-catalyzed alkyne–azide cycloaddition (CuAAC) reaction. The strands bound to biotin were isolated by binding to streptavidin magnetic beads followed by cleavage with hydrazine. Libraries were prepared, and sequences analyzed using paired-end 50 nucleotide reads. We obtained over 78 million reads for all samples, which were aligned to the reference human genome (hg38), and >94% reads were successfully mapped to the reference with < 5 % PCR duplicates. The genomic distribution of EBndG incorporation is shown in Figure 2a. The mapped reads were binned into 50 kb regions, normalized, and the difference between EBndG treated and untreated cells determined. As shown in Figure 2b and c, the 100 μM, but not 20 μM, concentration of EBndG produced regions that differed in the extent of EBndG incorporation.

**Figure 2.**
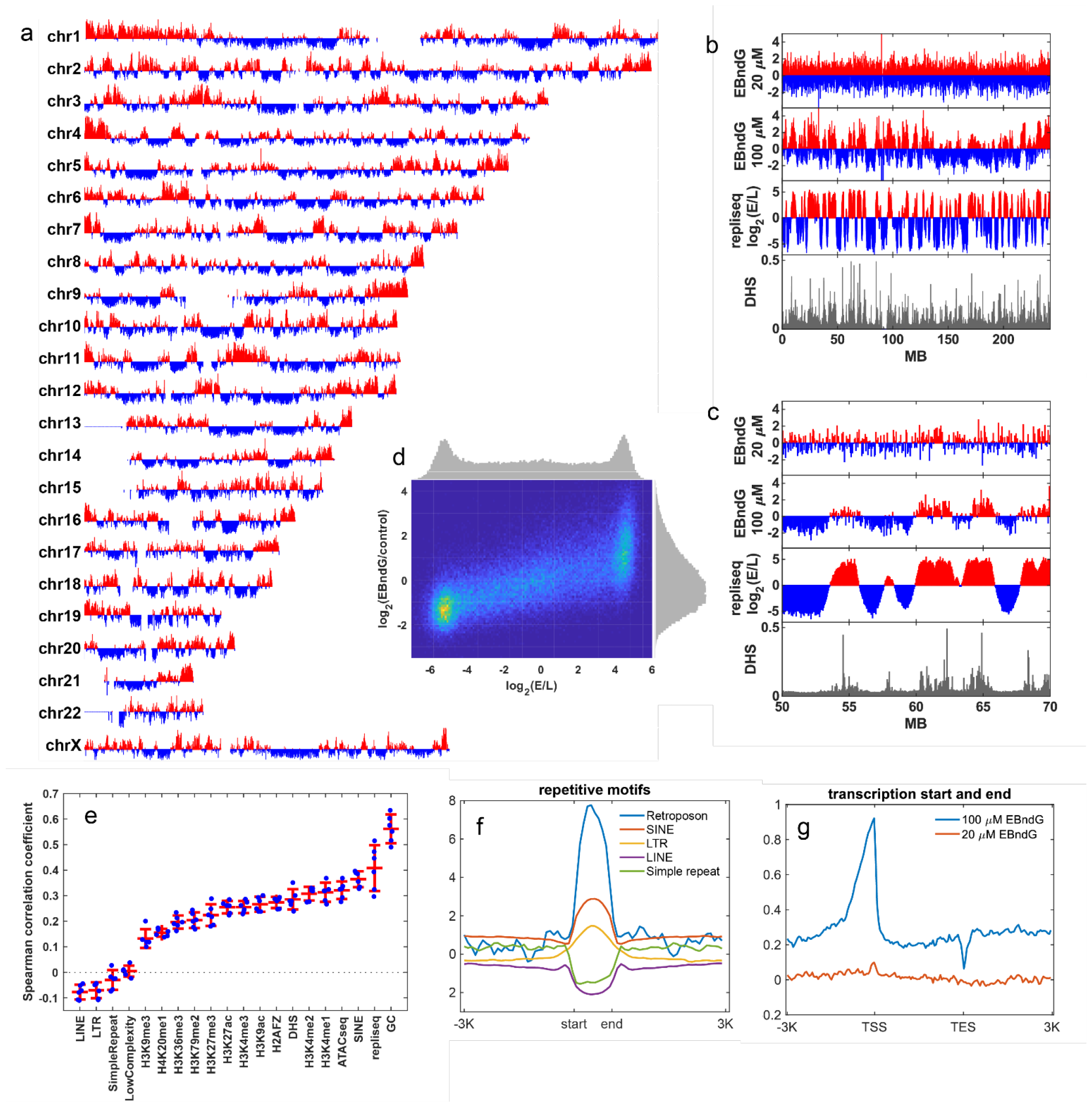
Distribution of EBndG in genomic DNA. (a) Genome-wide map of EBndG in cells exposed to 100 *μ*M EBndG. The function plotted is log_2_ (EBndG/control) calculated in 100 kb bins across six biological replicates. The control is untreated DNA (0 μM EBndG). (b) Detailed profile of chromosome 2 in which the EBndG distributed determined at 20 and 100 μM is compared with the replication timing score and the DNase hypersensitive sites. (c) Chromosome 2 is examined from 50 to 70 MB. (d) Two dimensional histogram in which the 100 μM EBndG abundance is compared to the replication timing score plotted at a 50kb bins.

### Comparisons of EBndG with genomic data

After determining the EBndG genome-wide distribution, we examined whether EBndG was accumulated or depleted in regions containing different genomic features. We compared our data with previously analyzed DNase hypersensitivity sites, histone epigenetic marks, various repetitive sequences and replication timing. The literature results were binned into 50 kb regions, and the Spearman’s correlation coefficients were obtained over the six EBndG biological replicates. The results are plotted in Figure 2e. The experiments with the highest correlation to the EBndG incorporation were the GC content, early replication timing, DNase hypersensitivity and ATAC-seq, regions that are associated with open DNA, as well as the SINE repetitive sequences. Small inverse correlations were associated with the LINE, LTP and low complexity repetitive sequences.

Histone modifications that are associated with active genes gave higher correlation coefficients than those associated with inactive chromatin. The heterochromatin-associated H3K9me3 had R = 0.13 ± 0.03 while H3K4me1, which is associated with primed enhancers, produced R= 0.31 ± 0.04, and H3K4me2, which his associated with active genes, had R= 0.31 ± 0.03. In general, the histone modification associated with active / open chromatin correlated to EBndG incorporation to a greater extent than those associated with heterochromatin. This trend was also observed with DNase hypersensitivity (DHS) regions and ATAC-seq that identify open regions of the DNA.

The two major repetitive sequences produced opposite correlations with EBndG, the short interspersed nuclear elements (SINEs, R = 0.36 ± 0.03) and long interspersed nuclear elements (LINEs) R = −0.08 ± 0.03). These correlations were observed in the sequence analysis in which the EBndG incorporation was compared with the start and end of the repetitive elements throughout the genome in Figure 2g. The regions associated with retroposons, SINE and LTSs had increased EBndG incorporation when compared with the surrounding DNA. The opposite was observed for LINEs and simple repeats. SINEs are associated with early replication timing while LINEs are associated with later replication (8) We also examiFigure 2ned EBndG incorporation against DNA replication timing in which replication score is log_2_(E/L) in which E/L is the ratio of BrdU incorporation in early and late Sphase. The incorporation of EBndG correlated with early replication times with R = 0.41 ± 0.09. This association was evident when examining replication timing versus EBndG incorporation in Figure 2b and c. Figure 2d shows a 2D histogram in which genomic EBndG and replication timing were compared in 50K bins throughout the genome (9). There is a clear relationship in which genomic regions with higher levels of EBndG were regions that were replicated early (upper right quadrant of Figure 2d).

To analyze the influence of gene transcription on EBndG incorporation, we examined the extent of incorporation over the transcription start and end sites in the genome. As shown in Figure 2f, EBndG incorporation was enriched in the promoter regions.

## Discussion

The present work describes the development of POLK-Seq, a methodology to identify the activity of pol κ in the genome. Central to this method is the design and synthesis of a dNTP substrate that has high selectivity for a single polymerase. Using mass spectrometry and flow cytometry, we found that EBndG was incorporated into the genomic DNA at low levels across all phases of the cell cycle (Figure 2c).

The genome-wide incorporation of EBndG was examined using whole genome sequencing. In general, the incorporation of EBndG was correlated with open regions of the genome as well as regions of the genome that are replicated early in S-phase. These results suggest that pol κ has an important cellular role that is distinct from translesion DNA synthesis during S-phase.

There have been many studies highlighting the importance of pol κ in S-phase especially in the bypass of DNA adducts and in the replication of repetitive sequences. These events typically occur in late S-phase due to several factors. Heterochromatin, which is typically replicated in late S phase, is also repaired less efficiently due to the absence of transcription coupled repair. In addition, the replication fork does not necessarily undergo the polymerase switch mechanism upon meeting a DNA adduct. Primpol is known to restart the fork downstream of the adduct, where upon the single-strand gap containing the DNA adduct is filled in in late Sphase.

Pol κ was shown to be important to nucleotide excision repair (4, 10) that occurs throughout the cell cycle. In addition, pol κ is recruited to UV damage by the Mettl3/Mettl14 *N*^6^-adenosine methyltransferase (11) involved transcriptional regulation (12). These events play roles throughout the cell cycle.

## Materials and Methods

### Reagents

AF488 picolyl azide and Dde biotin picolyl azide were purchased from Click Chemistry Tools (Scottsdale, AZ). DNA Isolation Midi-Prep Plus and DNA Clean and Concentrator Kits were purchased from Zymo Research. Fetal bovine serum (FBS) was purchased from Biowest (Bradenton, FL). Dynabeads MyOne Streptavidin C1 beads were purchased from ThermoFisher. *N*^2^-4-Ethynylbenzyl-2’-deoxyguanosine (EBndG) was synthesized as described.(13)

### Cell Culture and Treatment

GM12878 human lymphocytes (ATCC-NIST-8398), an ENCODE project Tier 1 human lymphocyte cell line (14), were cultured in Roswell Park Memorial Institute (RPMI) 1640 media supplemented with 15% FBS and 1% penicillin-streptomycin and 2 mM GlutaMax (Corning). The cells were grown at 37 °C with 5% CO_2_ at ∼80% relative humidity. They were seeded at 300,000 cells/mL and were split when they reached 10^6^ cells/mL. The day before experiments, cells were split to a concentration of 500,000 cells/mL in each flask. These cells were then treated with either 20 μM EdU for two hours or EBndG for four hours. Following this incubation, cells were harvested and washed with phosphate buffered saline (PBS) and pelleted. Cells used for DNA isolation were stored at −20°C until use.

### Flow Cytometry

The cells were treated with 50 μM EdU or 100 μM EBndG for various times, then harvested by centrifugation (5 min at 500×g). The cells were (a) washed with PBS, (b) treated with 4% paraformaldehyde in PBS for 15 min at room temperature, (c) washed with 1% bovine serum albumin (BSA) in PBS (w/v) and (d) treated with 1 mL 5% saponin in PBS (v/v) for 10 min. Following saponin treatment, the cells were incubated with the Click Reagent (1 mM CuSO_4_, 5 μM AF488 Picolyl Azide, 10 mM sodium ascorbate in PBS) for 30 min at room temperature. The cells were washed twice with 5% saponin in PBS (v/v) and treated with propidium iodide (PI)-RNase A reagent (75 μM PI, 10 μg/mL RNase A in 1% BSA/PBS). Finally, the cells were analyzed using a BD LSR Fortessa flow cytometer.

### DNA isolation

Treated cells (3 × 10^7^) were suspended in 20 mL 10 mM Tris-HCl, pH 7.4, 0.5 % SDS, 1 mM EDTA, 150 mM NaCl with 2 mg proteinase K and heated at 55° C for 2h. The solution was cooled and treated with one volume of phenol/CHCl_3_/iso-amyl alcohol (25/24/1). Following vigorous shaking the layers were separated by centrifugation. The aqueous layer was extracted an additional time and the DNA precipitated by addition of 2 mL 3 M sodium acetate and 15 mL isopropanol. The DNA was resuspended in 10 mL Tris-HCl, (pH 7.4) 1 mM EDTA and treated with 50 μg RNase A for 1 h at 37°C, followed by 100 μg proteinase K for 1 h at 55°C. The aqueous layer was extracted once with phenol/CHCl_3_/iso-amyl alcohol followed by a CHCl_3_ extraction. DNA was precipitated with 1/10 th volume 3 M sodium acetate and 2.5 volumes of ethanol.

### EBndG incorporation analysis by HPLC MS/MS

The DNA was redissolved in 10 mM Tris-HCl, pH 8, 10 mM MgCl_2_, 5 mM CaCl_2_, and mixed with DNase I (1 unit), phosphodiesterase I (0.005 units) and alkaline phosphatase (10 units) and incubated for 16 h at 37°C. After incubation, 25 μL of hydrolysate was taken for the analysis of dG by HPLC with UV detection. The remaining hydrolysate was loaded on a Strata-X cartridge (30 mg, Phenomenex) activated with 1 mL MeOH and 1 mL H_2_O. The cartridge was washed with 1 mL H_2_O, 1 mL of 20% MeOH, after which EBndG was eluted with 2 mL of 90% MeOH. The 90% MeOH fraction was concentrated to dryness in a rotary evaporator and redissolved in 50% MeOH. The samples were analyzed using a Sciex QTRAP 6500+ mass spectrometry coupled with a Sciex EXion UHPLC separation system. A 1.7 μm Acquity UPLC BEH C18 analytical column (2.1 × 100 mm, Waters, Ireland) was used to separate EBndG from other impurities. The mobile phase consisted of (A) 0.1 % acetic acid in water, and (B) acetonitrile. The gradient elution was conducted using a flow rate of 0.3 mL/min with a linear gradient from 10% B to 100% B over 3.6 min, followed by holding at 100% B for 1.4 min.

The Sciex QTrap 6500+ mass spectrometer was equipped with an electrospray ionization probe operated in positive mode. The decluster potential (DP) was 57 V, the entrance potential (EP) was 10 V, the collision energy (CE) was 24.7 V, and the collision cell exit potential (CXP) was 15 V, while the curtain gas (CUR) was 35 L/h, the collision gas (CAD) was medium, the ionspray voltage was 5000 V, the temperature was 250 °C, gas 1 was 20 L/h, and gas 2 was 20 L/h. The multiple reaction monitoring mode (MRM) was used to analyze and quantify EBndG, with the transitions of *m/z* 382 [M + H]^+^ > 266 [M – deoxyribose + H]^+^. The peak was integrated and quantified by Sciex OS 3.0 software.

At the dose and time point of our study, EBndG was incorporated at a level of 2.5 EBndG per million dG. This level is approximately 1/10th the level of *N*^2^-butyl-2’-deoxyguanosine observed in HEK cells (15).

### DNA fragmentation and NGS

DNA (20 μg) in 130 μL 10mM Tris-HCl pH 8.0 was fragmented using a Covaris E220 focused ultrasonicator in a microTUBE AFA Fiber Snap-Cap (Covaris, 520045) with the Duty Factor set to 10 and Peak Incident Power set to 175W at 200 cycles per burst for 180 seconds per sample. The average size of the DNA was 200 base pairs as determined by agarose gel electrophoresis.

The sheared DNA was reacted with 1mM CuSO_4_, 2mM THPTA, 40μM Dde biotin picolyl azide, and 5mM sodium ascorbate in PBS for 60 min with gentle rotation at room temperature. The DNA was separated from the biotin reagents with the Zymo DNA Clean and Concentrator-25 kit, with elution into 10 mM Tris-HCl (pH 8.0), 0.1 mM EDTA. Five percent of each DNA sample was kept aside as an input control.

MyOne Streptavidin C1 magnetic beads were used to capture the biotinated DNA and hydrazine was used to release the DNA. To prepare the beads for DNA binding, 10 μL of the DynaBeads were washed with Tween-20 (0.1 %) in PBS followed by 10 mM Tris-HCl (pH 8.0), 1 M NaCl twice. The beads were suspended in 50 μL 10 mM Tris, pH 8, 2 M NaCl followed by 50 μL of the DNA solution. The suspension was incubated for 1 h, washed twice with 10 mM Tris-HCl (pH 8.0), 1 M NaCl, followed by 10 mM Tris-HCl (pH 8.0) twice. The DNA was released from the beads with 50 μL 2% hydrazine (w/v) in 10 mM Tris-HCl (pH 8.0). The DNA was isolated with a Zymo DNA Clean and Concentrator-5 kit, with elution into 25 μL 10 mM Tris-HCl, pH 8. The DNA yield was 3-10 ng as determined by fluorometric analysis using a Qubit dsDNA high sensitivity assay kit.

The libraries were prepared using the NEB Ultra II DNA Library Prep Kit for Illumina. The samples were submitted to the Penn State College of Medicine Genome Sciences Facility. NGS was performed on a Novaseq Illumina instrument with a 50 cycle paired-end protocol.

### Data analysis

The FASTQ files were mapped onto the hg38 genome with Bowtie2 (16). Proper pairs were selected and duplicates were removed with Samtools (17). The reads were binned with Deeptools multiBamSummary (18). The data were normalized, the difference between the sample (EBndG = 20, 100 μM) and control (EBndG = 0 μM) obtained and plotted with Matlab.

We downloaded the call sets from the ENCODE portal (https://www.encodeproject.org/) (14) with the following identifiers ENCFF023LTU (H3K27ac), ENCFF131ZGZ (H3K79me2), ENCFF283LNH (H3K4me2), ENCFF291DHI (H3K27me3), ENCFF320OGZ (H3K4me3), ENCFF321BVG (H3K4me1), ENCFF377OJG (H2AFZ), (H4K20me1), ENCFF470YYO (ATAC-seq), ENCFF725UFY (H3K9me3), ENCFF759OLD (DNase-seq), ENCFF981JOU (H3K9ac).

The bed files for the repetitive sequences were extracted from 4DNI6. The data from replication timing was downloaded from 4D Nucleome Data Portal (https://data.4dnucleome.org/) (19) with the identifiers, 4DNFI6TILWWX, 4DNFIIMJQ8NT, 4DNFIS7J9B9X, 4DNFIT26294Y.

## Acknowledgments

This work was supported by grant number 5R01ES021762-10 from the NIH. This The Genome Sciences Core (RRID:SCR_021123), Flow Cytometry Core (RRID:SCR_021134) and the Mass Spectrometry and Proteomics Core (RRID:SCR_017831) services and instruments used in this project were funded, in part, by the Pennsylvania State University College of Medicine and the Pennsylvania Department of Health using Tobacco Settlement Funds (CURE).

